# Fine-tuning quantitative agronomic traits by manipulating gene copy number in rice

**DOI:** 10.1101/2025.05.16.654416

**Authors:** Chihiro Nomura, Hiroyuki Kanzaki, Eiko Kanzaki, Motoki Shimizu, Kaori Oikawa, Hiroe Utsushi, Kazue Ito, Yusaku Sugimura, Ryohei Terauchi, Akira Abe

## Abstract

Although recent plant pan-genome studies have revealed extensive copy number variations, there are limited studies addressing the phenotypic consequences of these variations. Here, we manipulated the copy number of OsMADS18 in rice (Oryza sativa) through gene editing and evaluated the effects. OsMADS18, which encodes a MADS-box transcription factor known to regulate plant development, is tandemly duplicated in the elite rice cultivar ‘Hitomebore’ as a natural variant. We therefore edited the OsMADS18 copy number using a basic CRISPR/Cas9 system and established rice lines harboring one to three tandem copies of OsMADS18, as identified through high throughput qPCR screening. The presence of one to three OsMADS18 tandem copies was reflected in stepwise increases in transcript levels and concomitant trait values for leaf blade length and culm length. These results demonstrate that manipulating gene copy number can fine-tune important agronomic traits, providing a novel breeding strategy for crop improvement.

## Introduction

Since the advent of high-throughput sequencing, numerous single-nucleotide polymorphisms (SNPs) and short insertion/deletion polymorphisms (indels) have been identified that affect phenotypes in many organisms, from humans (Koboldt *et al*., 2013) to crop plants (Singh *et al*., 2022). Recent developments in long-read sequencing technologies, such as the PacBio and Oxford Nanopore Technologies platforms, have helped reveal copy number variations (CNVs) and large structural variations in the genome, which were previously difficult to capture using short-read sequencing methods, such as those involving the Illumina platform (Alonge *et al*., 2020; Liu *et al*., 2020; Qin *et al*., 2021). However, current *in silico* analyses exploring the relationships between CNV and phenotype using genome sequences obtained through long-read sequencing among accessions of a single species are based on materials with diverse genetic backgrounds, making it difficult to demonstrate a direct dosage effect of CNV (Yuan *et al*., 2021). Therefore, there is a pressing need to study how variation in gene dosage caused by CNV results in different transcript levels and ultimately influences plant phenotypes (Maino *et al*., 2021).

Genome editing is a powerful tool for introducing mutations into target genomic locations (Gaj *et al*., 2013). Among gene-editing methods, the clustered regularly interspaced short palindromic repeats (CRISPR)/CRISPR-associated nuclease 9 (Cas9) system is the most widely used for genetic research (Jinek *et al*., 2012; Doudna & Charpentier, 2014; Belhaj *et al*., 2015). CRISPR/Cas9 typically cuts the genome near a target site bound by a single guide RNA (sgRNA), after which the repair machinery introduces mutations such as indels. CRISPR/Cas9-mediated gene editing can be used to knock out genes to explore gene function (Shalem *et al*., 2015; Bao *et al*., 2019). However, a complete loss-of-function mutant often displays drastic changes in phenotype compared to the wild type (Pnueli *et al*., 1998; Jeon *et al*., 2000; Komatsu *et al*., 2003; Qi *et al*., 2008) and thus is not suitable for studying quantitative phenotypic effects associated with subtle changes in gene expression. In addition to knocking out gene function, CRISPR/Cas9-mediated genome editing can also delete larger genomic fragments by using two sgRNAs that target two sites flanking the genomic sequence to be deleted (Belhaj *et al*., 2015).

We hypothesized that this approach could be used to increase or decrease the copy number of a target gene, thereby revealing the quantitative effects of gene CNV on expression levels and phenotypes. In this study, we manipulated the copy number of the MADS-box transcription factor gene *OsMADS18* in rice (*Oryza sativa*) through gene editing and evaluated the effects, providing a powerful approach for fine-tuning complex traits in crops.

## Materials and Methods

### Plant Materials and Growth Conditions

The genomic structure of the target *OsMADS18* region was compared between rice (*Oryza sativa*) ssp. *japonica* cultivars Hitomebore and Sasanishiki. Genome editing was performed using Hitomebore, which carries two tandem copies of *OsMADS18*. Plants with *OsMADS18* CNVs were selected from 107 regenerated gene-edited independent lines. The selected individuals were allowed to self-pollinate to establish homozygous lines with different *OsMADS18* copy numbers. These lines were grown in a plant incubator to the seedling stage. After transplanting, the plants were grown in a temperature-controlled greenhouse under short-day conditions, with day/night temperatures of ∼27°C/∼23°C. The *OsMADS18* transcript levels and various phenotypes were evaluated.

### Plasmid Construction and Plant Transformation

To generate lines with CNV at *OsMADS18*, one sgRNA targeting the duplicated genomic region was designed using CRISPRdirect (https://crispr.dbcls.jp/) (Naito *et al*., 2015). To increase the probability of mutations, a binary vector containing two sgRNA expression cassettes was constructed based on a previous study (Ma *et al*., 2015). The *OsU6a* and *OsU6b* promoters, amplified using pYLsgRNA-OsU6a and pYLsgRNA-OsU6b as templates, were each fused to the target sequence and sgRNA scaffold. The ccdB cassette of the pYLCRISPR/Cas9P_ubi_-H vector was replaced with the two sgRNA cassettes via the BsaI site using Golden Gate assembly. This vector was introduced into *Agrobacterium tumefaciens* strain EHA105, and rice transformation in the Hitomebore background was performed as described previously (Toki *et al*., 2006). The pYLCRISPR/Cas9P_ubi_-H (RRID:Addgene_66187), pYLsgRNA-OsU6a (RRID:Addgene_66194), and pYLsgRNA-OsU6b (RRID:Addgene_66196) vectors were kindly provided by Yao-Guang Liu. The primers used in this study are listed in Table **S1**.

### *De Novo* Assembly

To reconstruct the Hitomebore and Sasanishiki genomes, *de novo* assembly of each genome was performed using Nanopore long reads and Illumina short reads following a previously published method (Sugihara *et al*., 2023). To extract high-molecular-weight genomic DNA from rice leaf tissue for Nanopore sequencing, leaf blades were carefully ground using a mortar and pestle while being cooled with liquid nitrogen. Genomic DNA was then extracted using a NucleoBond High-molecular-weight DNA Kit (Macherey-Nagel, Düren, Nordrhein-Westfalen, Germany). Following DNA extraction, low-molecular-weight DNA was eliminated using a Short Read Eliminator Kit XL (Circulomics, Baltimore, MD, USA). For Hitomebore, genomic library preparation was performed using a Ligation Sequencing Kit (SQK-LSK-114; Oxford Nanopore Technologies [ONT], Oxford, Oxfordshire, UK) according to the manufacturer’s instructions, and sequencing was performed using PromethION 2 Solo (ONT). Long reads for Hitomebore produced on the PromethION (ONT) platform, as described by Sugihara *et al*. (2023), were also used. For Sasanishiki, genomic library preparation was performed using a Ligation Sequencing Kit (SQK-LSK-109; ONT), and sequencing was performed on a MinION instrument.

Base-calling of the Nanopore long reads was conducted using Dorado v0.9.0 (https://github.com/nanoporetech/dorado). The first 50 bp of each read and reads with an average read quality score below 10 were removed using Chopper v0.9.0 (De Coster & Rademakers, 2022). The resulting clean Nanopore long reads were assembled using NECAT v0.0.1 (Chen *et al*., 2021), with the genome size set to 380 Mb and with reads longer than 3,000 bp. To improve the accuracy of the assembly, Racon v1.4.20 (Vaser *et al*., 2017) was employed for error correction using Nanopore reads, followed by Medaka v1.4.1 (https://github.com/nanoporetech/medaka) for misassembly correction. Two rounds of consensus correction were then performed using BWA v0.7.17 (Li, 2013) and HyPo v1.0.3 (Kundu *et al*., 2019) with the Illumina short reads. Redundant contigs were removed using Purge Haplotigs v1.1.1 (Roach *et al*., 2018), resulting in a *de novo* assembly of 376.2 Mb comprising 59 contigs for Hitomebore and 378.7 Mb consisting of 49 contigs for Sasanishiki.

*De novo* assembly was also performed for Hitomebore gene-edited lines with one or three copies of the *OsMADS18* region. In both cases, high-molecular-weight genomic DNA was extracted as described above. The library for the single-copy line was prepared using an SQK-LSK110 Kit and sequenced on a MinION instrument, while the library for the three-copy line was prepared using an SQK-LSK114 Kit and sequenced on a PromethION 2 Solo instrument. Base-calling was performed as described above, and *de novo* assembly was conducted using Flye v2.9.5 (Kolmogorov *et al*., 2019).

### Illumina-Based DNA Sequencing

Total genomic DNA was extracted from rice leaf blades using the CTAB-based method with a NucleoSpin Plant II Kit (Macherey-Nagel). Sequencing libraries were prepared using a Collibri ES DNA Library Prep Kit for Illumina Systems (Thermo Fisher Scientific, Waltham, MA, USA). Libraries were sequenced on a MiSeq (Illumina, San Diego, CA, USA) instrument in 251-bp paired-end mode for Hitomebore and in 301-bp paired-end mode for Sasanishiki and on a NovaSeq X Plus (Illumina) instrument in 151-bp paired-end mode for lines with CNV (ranging from one to three copies) at *OsMADS18*. The resulting raw reads from the MiSeq instrument were filtered using Trimmomatic v0.36 (Bolger *et al*., 2014) with the paired-end option. Adapter sequences were removed using the ILLUMINACLIP:TruSeq3-PE-2.fa:2:30:10 option. Bases with quality scores of <20 at the beginning and end of reads were removed (LEADING:20, TRAILING:20). A sliding window of four bases was used to remove reads with average quality scores of <20 (SLIDINGWINDOW:4:20). Reads were removed if the read length after trimming was <36 (MINLEN:36) and were cut to 200 bases if the read length was >200 bases (CROP:200). The resulting raw reads from the NovaSeq X Plus instrument were filtered using fastp v0.22.0 (Chen *et al*., 2018) with default settings supplemented with the -- detect_adapter_for_pe option to automatically detect and trim adapter sequences and the --trim_poly_g option to trim polyG tails. The processed reads were aligned to the rice reference genome IRGSP-1.0 using BWA v0.7.18. The resulting SAM files were converted to BAM format, sorted, and indexed using SAMtools v1.9 (Danecek *et al*., 2021).

### Analysis of Sequencing Depth

Sequencing depth was calculated using mosdepth v0.3.3 (Pedersen & Quinlan, 2017) with the BAM files generated from high-throughput sequencing and subsequent processing. The sliding window used in the analysis was created with BEDTools v2.31.1 (Quinlan, 2014) using the command ‘makewindows’. Relative sequencing depth between the target and control was calculated using the following equation:

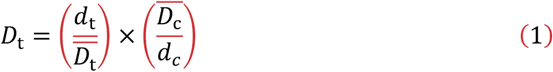

where *D*_t_ is the relative sequencing depth in the region of the target genome, *d*_*t*_ is the sequencing depth in the region of the target genome, 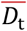 is the average sequencing depth of the whole target genome, *d*_c_ is the sequencing depth in the target region of the control genome, and 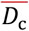 is the average sequencing depth of the whole control genome.

### Comparison of Genome Structures

The genomic structure of the target *OsMADS18* region was compared using MUMmer4 v4.0.1 (Marçais *et al*., 2018). First, the ‘blastn’ command from BLAST+ v2.13.0 (Camacho *et al*., 2009) was used to identify the contig containing *OsMADS18* from each of the four *de novo* assemblies. The identified contigs were then extracted individually using SeqKit v2.4.0 (Shen *et al*., 2024). Each contig was compared to chromosome 7 of the IRGSP-1.0 reference genome using the ‘nucmer’ command of MUMmer4. The results were filtered using the ‘delta-filter’ command, setting the minimum alignment identity to 95% and the minimum alignment length to 5,000 bp. Finally, the filtered results were visualized using the ‘mummerplot’ command.

To identify SNPs and indels within the duplicated regions of the Hitomebore genome, the duplicated sequences were separated into two segments corresponding to the upstream and downstream copies. These sequences and the corresponding region from the IRGSP-1.0 reference genome were aligned using MAFFT v7 (Katoh & Standley, 2013). Sequence differences were then identified based on the alignment results.

The duplicated regions in Hitomebore and the corresponding region in Sasanishiki were visualized using the Integrative Genomics Viewer (IGV) v2.16.2 (Robinson *et al*., 2011), with the IRGSP-1.0 reference genome.

### qPCR-Based Analysis of Target Gene Copy Number

qPCR was conducted using a QuantStudio 3 Real-Time PCR System (Thermo Fisher Scientific) and Luna Universal qPCR Master Mix (New England Biolabs, Ipswich, MA, USA). To estimate the CNV of *OsMADS18*, the relative amplification ratio was calculated using qPCR data from primer sets designed to span the duplicated junction and a non-duplicated control region. The analysis was based on the Pfaffl approach (Pfaffl, 2001), which incorporates primer-specific amplification efficiencies. The mean *C*_t_ value of wild-type Hitomebore plants (homozygous for two copies of *OsMADS18, n* = 3) was used as the calibrator. The *C*_t_ difference (Δ *C*_t_) for each primer set was calculated using Equations 2 and 3:

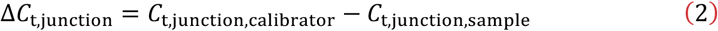

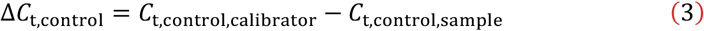

The relative amplification ratio (*R*) was then calculated using Equation 4:

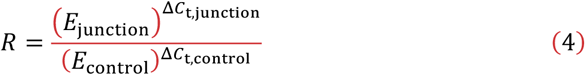

where *E*_junction_ and *E*_control_ represent the amplification efficiencies of the junction and control primer sets, respectively.

Further details are provided in Fig. **S1**, and the primer sequences used in the experiments are listed in Table **S1**.

### RT-qPCR Analysis

Leaf blade samples were bulk collected from the uppermost fully expanded leaf of the main stem, with three individuals per sample, and immediately frozen in liquid nitrogen. Each sample was ground to a powder, and total RNA was extracted from the samples using a NucleoSpin RNA Plant Kit (Macherey-Nagel). The total RNA was reverse transcribed into first-strand cDNA using PrimeScript RT Master Mix (Takara Bio, Kusatsu, Shiga, Japan). qPCR was performed using a QuantStudio 3 Real-Time PCR System with Luna Universal qPCR Master Mix. The ΔΔ*C*_t_ method was used to quantify transcript levels, and the relative transcript levels of *OsMADS18* were calculated using *UBQ5* (Jain *et al*., 2006, 2018) as the internal control. The mean *C*_t_ value of biological replicates from plants with one homozygous copy of *OsMADS18* was used as the calibrator. The primers used in the experiment are listed in Table **S1**.

### Phenotyping

After heading, fully expanded leaf blades were collected, scanned, and analyzed using a custom-developed automated measurement program in Python to calculate their morphology, including length. Leaf blade length was compared among individuals with the same total number of leaves on the main stem. After heading was fully completed, the length from the ground level to the panicle neck node was measured and defined as culm length. The number of days from sowing to heading was recorded as the heading date. All traits were measured on the main stem.

## Statistical Analysis

Trait comparisons were made between lines with different *OsMADS18* copy numbers, and statistical significance was determined using the Tukey-Kramer test in R v.4.4.2 (https://www.r-project.org/). Details of the test are provided in the figure legends.

## Results

### *OsMADS18* is Tandemly Duplicated in Hitomebore

MADS-box transcription factors are widely distributed across eukaryotes. For example, OsMADS18 from rice is involved in the transition from the vegetative to the reproductive stages (Fornara *et al*., 2004; Kobayashi *et al*., 2012) and exerts pleiotropic effects on germination and tillering (Yin *et al*., 2019). A previous study of a pan-genome from 33 genetically diverse rice accessions revealed CNVs for *OsMADS18* that arose via tandem duplication (Qin *et al*., 2021). Here, we generated *de novo* genome assemblies for the elite *japonica* rice cultivar ‘Hitomebore’ and the *japonica* cultivar ‘Sasanishiki’ using long reads generated by Oxford Nanopore sequencing and compared the results to the reference genome IRGSP-1.0 from the rice cultivar ‘Nipponbare’ (Kawahara *et al*., 2013) (Fig. **S2a,b**). We also assessed sequencing coverage using Illumina-based short reads between Hitomebore and Sasanishiki (Figs. **S2c, S3a,b**).

We confirmed the presence of a 20-kb tandem duplication at 24.776–24.796 Mb on chromosome 7 in the Hitomebore genome and its absence from the Sasanishiki and IRGSP-1.0 reference genomes. According to the Rice Annotation Project Database (RAP-DB) (Kawahara *et al*., 2013; Sakai *et al*., 2013), the only full-length protein-coding gene contained in this 20-kb fragment is *OsMADS18*, which is duplicated specifically in Hitomebore (Figs. **S2a–c, S3**). We detected only one SNP and three indels across the two tandem repeats of the 20-kb fragment, with no variation in the two copies of the *OsMADS18* coding region (Fig. **S2d**). Given that *OsMADS18* is the only full-length protein-coding gene within the duplicated region and that sequence variation between the two tandem copies is minimal, we reasoned that CNV of *OsMADS18* might provide an attractive model for the genetic evaluation of gene dosage effects. We thus attempted to locally manipulate the copy number of *OsMADS18* using CRISPR/Cas9 and study how changes in gene copy number would affect plant phenotypes.

### CRISPR-Mediated Manipulation of *OsMADS18* Copy Number

To manipulate the *OsMADS18* copy number, we designed an sgRNA targeting a site 12 kb upstream of the ATG of *OsMADS18*, within the region tandemly duplicated in Hitomebore, for CRISPR/Cas9-mediated editing in this cultivar. Since the two copies of *OsMADS18* have the same sequence, this sgRNA has two target sites in this cultivar (Fig. **1a**). Among the T_0_ individuals regenerated from Hitomebore calli, we looked for individuals with altered *OsMADS18* copy numbers using quantitative PCR (qPCR) with two primer sets (Fig. **S1**): one set amplifying an amplicon outside of the duplicated region, thus maintaining a constant copy number, and another set designed to anneal to either side of the junction of the duplicated region to specifically amplify the junction in the presence of a tandem duplication. We calculated the relative amplification ratio of these two primer sets using the efficiency-corrected Pfaffl method (Pfaffl, 2001). When the relative amplification ratio of Hitomebore (with two homozygous copies of the duplication) was set to 1, deviations of the relative ratio from 1 in increments or decrements of 0.5 units reflected a change in *OsMADS18* copy number (Fig. **S1**).

**Fig. 1.**
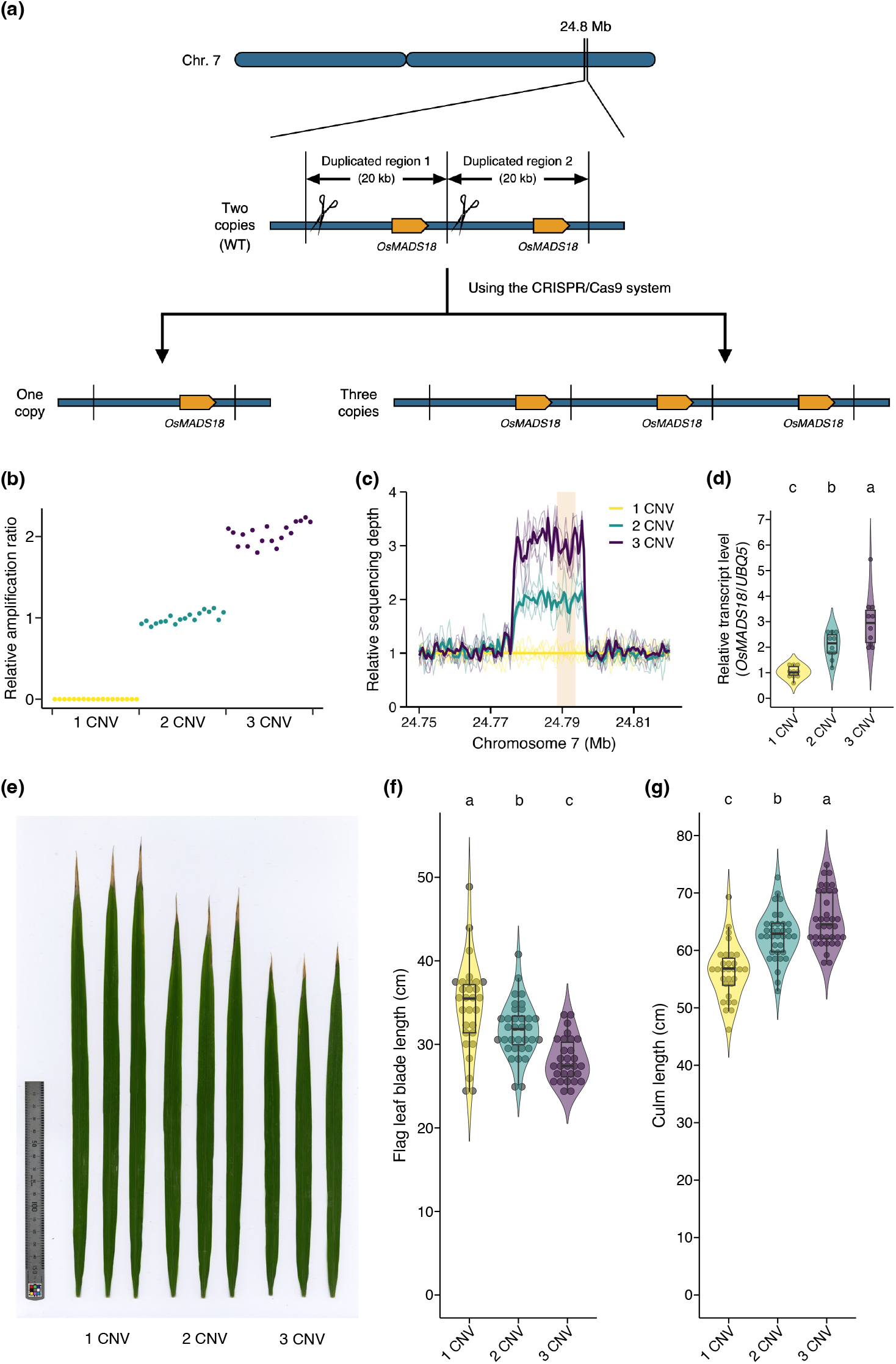
Phenotypic variation associated with the manipulation of *OsMADS18* copy number. (a) Diagram of the method used to induce CNV of *OsMADS18*. The sgRNA targets two identical sequences within the tandemly duplicated *OsMADS18* genomic region. CRISPR/Cas9-mediated editing can result in plants with either one or three copies of *OsMADS18*. Target sites of the sgRNA are indicated by scissor icons. (b) Relative amplification ratio of the tandemly duplicated *OsMADS18* region, as determined by qPCR (*n* = 18 for each CNV). (c) Relative sequencing depth of Illumina short reads around *OsMADS18*. A sliding window with a window size of 1,000 bp and a step size of 500 bp was used. Thin lines represent individual plants, and the thick lines indicate the mean depth for plants with each *OsMADS18* CNV (*n* = 5). The mean value for plants with one homozygous copy of *OsMADS18* was set to 1. The orange shaded area marks the position of *OsMADS18*. (d) Relative *OsMADS18* transcript levels in leaf blades, as determined by RT-qPCR and normalized to *UBQ5*, with the mean transcript level of plants with one copy set to 1. Leaf blades were bulk collected from three individuals at 58 days after sowing (*n* = 10; total of 30 plants per *OsMAD18* CNV). (e) Representative photograph of flag leaves. (f, g) Lengths of the flag leaf blade (*n* = 29, 34, and 29 for 1, 2, and 3 CNV, respectively) (f) and culm (*n* = 30, 35, and 36 for 1, 2, and 3 CNV, respectively) (g). “1 CNV,” “2 CNV,” and “3 CNV” refer to plants with one, two, and three homozygous copies of *OsMADS18*, respectively, for (b–g). Different lowercase letters indicate significant differences (*P* < 0.05) based on the Tukey-Kramer test for panels (d), (f), and (g). All violin plots were generated using kernel density estimation with a smoothing adjustment factor of 1.5. All internal boxplots represent the first and third quartiles (the lower and upper hinges, respectively), the median (the middle horizontal line), and the range from the hinges to the smallest and largest observations within 1.5 times the interquartile range (the bottom and top whiskers, respectively). All data points, including outliers in the boxplot, are displayed as dots in the dot plot.

Of the 107 regenerated T_0_ plants obtained from different calli or from the same callus with different culture periods, qPCR analysis identified nine plants with a relative amplification ratio of ∼0.5 and one with a relative amplification ratio of ∼1.5, suggesting the loss or gain of one *OsMADS18* copy, respectively (Fig. **S4a,b**; Dataset **S1**). We allowed these plants to self-pollinate and repeated the qPCR analysis on their progeny. From the parents with a relative amplification ratio of ∼0.5, we obtained progeny with relative amplification ratios of 0 (reflecting absence of the junction sequence of the duplicated region), ∼0.5, or ∼1 (Fig. **S4c**; Dataset **S1**). From the parents with a relative amplification ratio of ∼1.5, we obtained progeny with relative amplification ratios of ∼1, ∼1.5, or ∼2 (Fig. **S4d**; Dataset **S1**). Furthermore, plant lines with relative amplification ratios of ∼0, ∼1, or ∼2 produced progeny with the same relative amplification ratios, suggesting that these plants were homozygous for one, two, or three copies of *OsMADS18*, respectively (Fig. **1b**; Dataset **S2**). These results demonstrate that our qPCR-based approach is practical for efficiently detecting genomic CNVs.

To confirm these copy numbers, we examined the sequencing coverage depth over the *OsMADS18* genomic region using Illumina short reads from homozygous plants with an estimated one, two, or three tandem copies. The relative sequencing depth within the *OsMADS18* region was lower in plants with one copy of *OsMADS18* and higher in plants with three copies of *OsMADS18* relative to those with two copies of this gene (Fig. **1c**). *De novo* genome assembly using Nanopore long reads for the genomes of homozygous individuals with an estimated one or three *OsMADS18* copies in the Hitomebore background showed that the *OsMADS18* copy number was indeed one and three, respectively, as evidenced by dot matrix analysis (Fig. **S5**).

### Copy Number Variation at *OsMADS18* Modulates Agronomic Traits in a Stepwise Manner

To study the effect of CNV at *OsMADS18* on its transcript levels, we performed reverse-transcription qPCR (RT-qPCR) on individuals with one to three copies of the gene, normalizing to *UBQ5* (Jain *et al*., 2006, 2018), with the mean relative *OsMADS18* transcript level for plants with one copy set to 1. The relative *OsMADS18* transcript levels gradually increased from an average of 1 to 2 or 3 in plants homozygous for one, two, or three copies of *OsMADS18*, respectively, thus neatly correlating with gene copy number (Fig. **1d**; Dataset **S2**). These findings suggest that simple genome editing introduced CNV that resulted in stepwise changes in *OsMADS18* expression levels.

We assessed the agronomic traits of lines with CNV (from one to three copies) at *OsMADS18*. We observed no major difference in overall plant architecture among the rice lines (Fig. **S6a**). However, the flag leaf blade was significantly shorter when the *OsMADS18* copy number was higher (Fig. **1e,f**; Dataset **S2**), as was the length of the leaf blade just below the flag leaf (Fig. **S6b**; Dataset **S3**). By contrast, culm length was significantly greater in plants with more *OsMADS18* copies (Fig. **1g**; Dataset **S2**). To our knowledge, effects of *OsMADS18* on leaf blade size and culm length have not been previously reported. Previous studies have shown that *OsMADS18* is upregulated during the transition from vegetative to reproductive growth and is involved in the number of days to heading (Fornara *et al*., 2004; Kobayashi *et al*., 2012). However, in the current study, we measured no significant difference in the number of days to heading among the three sets of lines with CNV at *OsMADS18* (Fig. **S6c**; Dataset **S3**).

## Discussion

Most agronomic traits are quantitative in nature, and their variations are controlled by subtle differences in the expression levels of multiple genes. Thus, manipulating gene expression is an important method for adjusting traits of interest. Promoter editing was recently developed as a means to generate a large allelic series of plants with different expression levels of a target gene by inserting random mutations in the *cis*-regulatory elements of promoters (Rodríguez-Leal *et al*., 2017). However, because this approach targets *cis*-regulatory elements, it is difficult to predict the direction and degree of changes in expression levels in the edited plants. Moreover, the spatiotemporal gene expression pattern may differ from that driven by the intact promoter. We propose that manipulating gene copy number is a powerful alternative method to create lines with a range of copy numbers, leading to stepwise changes in the expression levels of a target gene without randomly perturbing its spatiotemporal expression patterns. This method should allow for the fine-tuning of important traits by optimizing gene copy number, thus accelerating plant breeding.

## Acknowledgements

We thank Yao-Guang Liu for providing the pYLCRISPR/Cas9P_ubi_-H, pYLsgRNA-OsU6a, and pYLsgRNA-OsU6b plasmids, as well as Yumiko Ogasawara and Miyuki Saito for technical support. Part of the analysis was performed using the National Institute of Genetics (NIG) supercomputer at the ROIS National Institute of Genetics. This research was partially supported by the following grants: the Research Program on the Development of Innovative Technology (JPJ007097) from the Project of the Bio-oriented Technology Research Advancement Institution (BRAIN); and JSPS KAKENHI (JP22K20584 to C.N. and JP23K26882 to A.A.).

## Competing interest

None declared.

## Author contributions

CN and AA conceived and designed the research. CN, HK, EK, MS, KO, HU, KI, YS, and AA performed the experiments. CN and AA conducted bioinformatics analyses. CN, RT, and AA wrote and revised the manuscript. All authors read and approved the final version of the manuscript.

## Data availability

The raw sequencing reads have been deposited in the DNA Data Bank of Japan (DDBJ) under BioProject IDs PRJDB13864 and PRJDB20676, with the accession numbers provided in Table S2. Nanopore-based *de novo* genome assemblies for four samples are available at Zenodo (DOI: https://doi.org/10.5281/zenodo.15335260). The IRGSP-1.0 reference genome used in this study was obtained from RAP-DB (https://rapdb.dna.affrc.go.jp/download/irgsp1.html). Source data used in this study are provided in Datasets S1, S2, and S3.

## Supporting information

**Dataset S1** Source data for all panels in Fig. S4.

**Dataset S2** Source data for panels (b), (d), (f), and (g) in Fig. 1.

**Dataset S3** Source data for panels (b) and (c) in Fig. S6.

**Fig. S1** Diagram of the method used to determine *OsMADS18* copy number by qPCR. **Fig. S2** Tandem duplication of the *OsMADS18* genomic region on chromosome 7 in Hitomebore.

**Fig. S3** Genome browser views of the *OsMADS18* genomic region on chromosome 7.

**Fig. S4** *OsMADS18* CNV in T_0_ plants and progeny as determined by qPCR.

**Fig. S5** Genomic structure of the *OsMADS18* locus in IRGSP-1.0 and CNV-edited lines.

**Fig. S6** Effects of *OsMADS18* CNV on other agronomic traits.

**Table S1** List of primers used in this study.

**Table S2** List of sequencing data and corresponding Sequence Read Archive (SRA) accession numbers.

## Notes

https://doi.org/10.5281/zenodo.15335260

### Competing Interest Statement

The authors have declared no competing interest.

### Summary of Updates

We made minor revisions to improve clarity and presentation.

