## Supplemental information for "Fine-tuning quantitative agronomic traits by manipulating gene copy number in rice"

The following Supporting Information is available for this article:

**Fig. S1** Diagram of the method used to determine *OsMADS18* copy number by qPCR.

**Fig. S2** Tandem duplication of the *OsMADS18* genomic region on chromosome 7 in Hitomebore.

**Fig. S3** Genome browser views of the *OsMADS18* genomic region on chromosome 7.

**Fig. S4** *OsMADS18* CNV in $T_0$ plants and progeny as determined by qPCR.

**Fig. S5** Genomic structure of the *OsMADS18* locus in IRGSP-1.0 and CNV-edited lines.

**Fig. S6** Effects of *OsMADS18* CNV on other agronomic traits.

**Table S1** List of primers used in this study.

**Table S2** List of sequencing data and corresponding Sequence Read Archive (SRA) accession numbers.

**Fig. S1** Diagram of the method used to determine *OsMADS18* copy number by qPCR. When the relative amplification ratio of Hitomebore (homozygous for two copies of *OsMADS18*) is set to 1, the values for different genotypes are as follows: homozygous for one copy, 0; heterozygous for one and two copies, 0.5; heterozygous for two and three copies, 1.5; homozygous for three copies, 2.

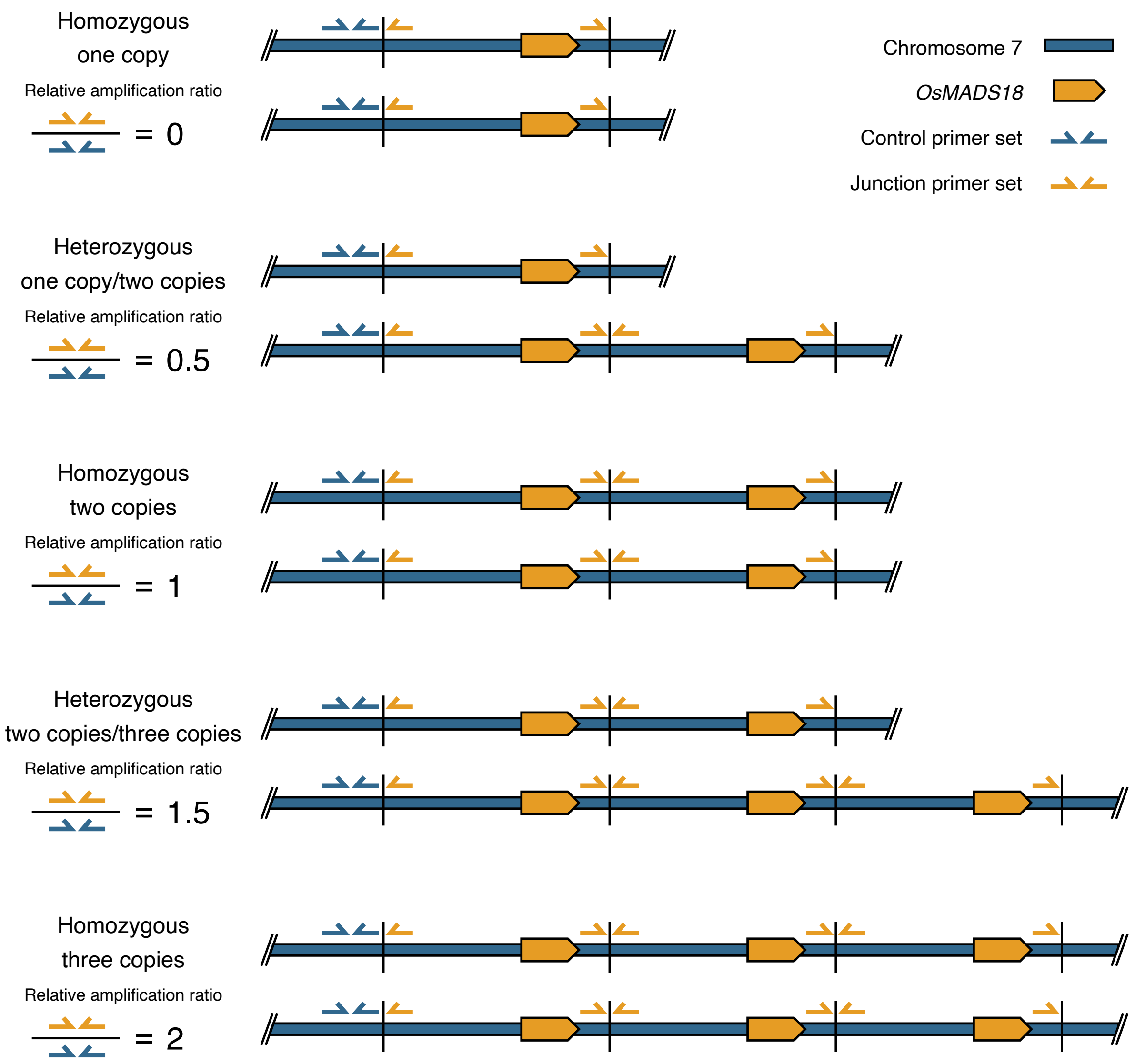

**Fig. S2** Tandem duplication of the *OsMADS18* genomic region on chromosome 7 in Hitomebore. (a, b) Dot matrix plots of the *OsMADS18* genomic region between Hitomebore and IRGSP-1.0 (a) and between Sasanishiki and IRGSP-1.0 (b). Grid lines are spaced at 0.03-Mb intervals. (c) Relative sequencing depth of Hitomebore (the target) to Sasanishiki (the control) using Illumina short reads around *OsMADS18*. A sliding window with a window size of 5,000 bp and a step size of 500 bp was used. The orange shaded area marks the position of *OsMADS18* for (a–c). (d) One SNP and three indels were observed in duplicated region 2 relative to duplicated region 1.

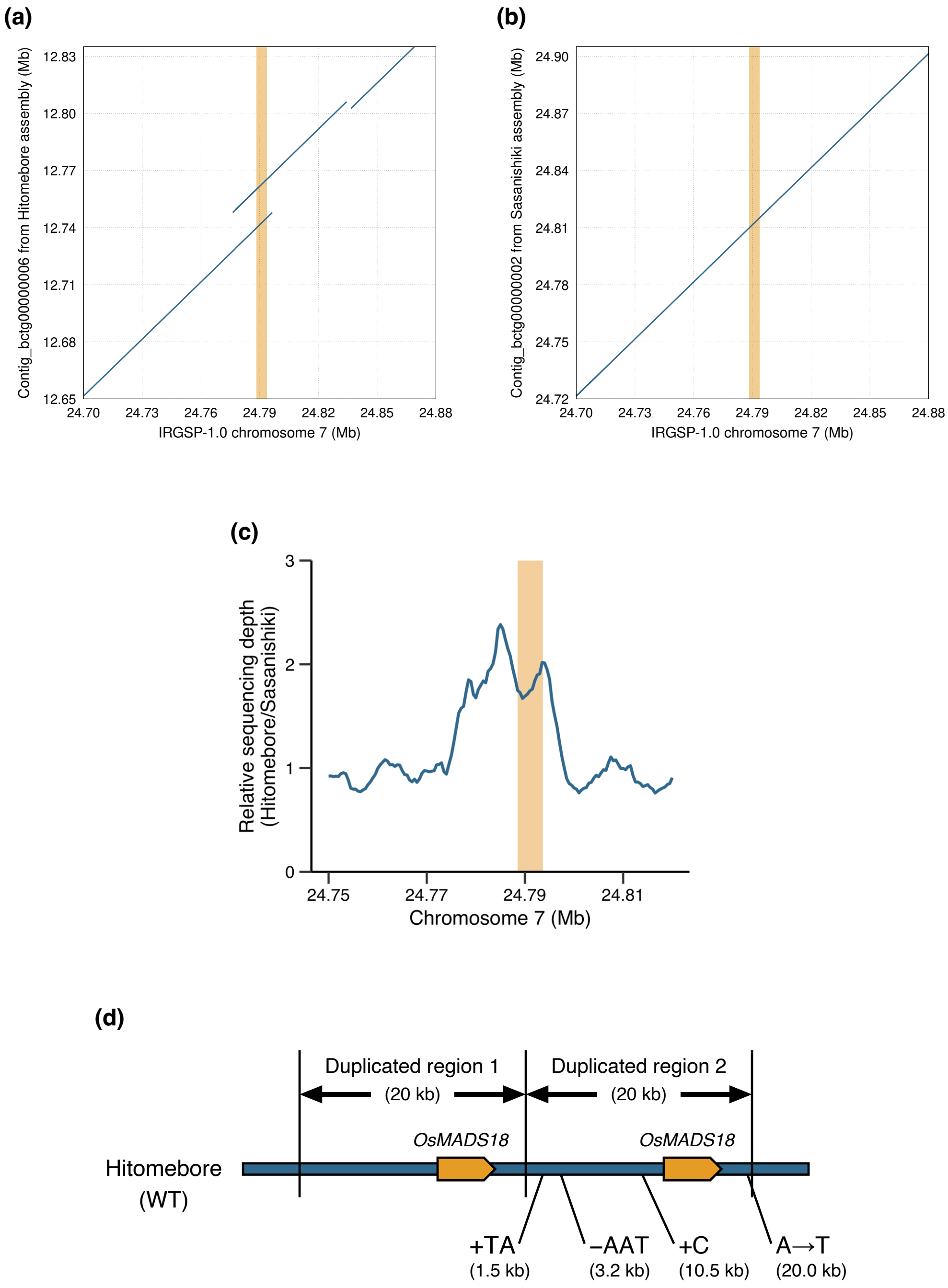

**Fig. S3** Genome browser views of the *OsMADS18* genomic region on chromosome 7. (a) IGV view of a 30-kb window in Hitomebore. The red horizontal bar indicates the duplicated segment. (b) IGV view of a 30-kb window in Sasanishiki. (c) JBrowse view of the duplicated *OsMADS18* region in RAP-DB.

**(a)**

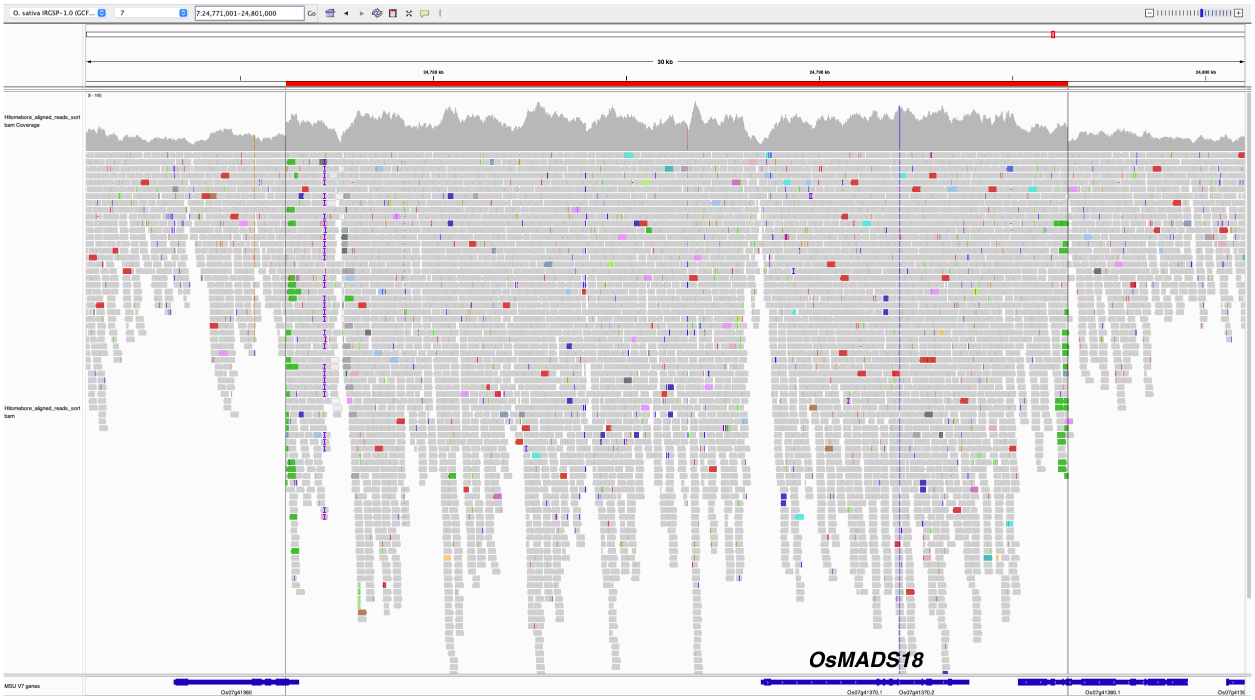


**(b)**

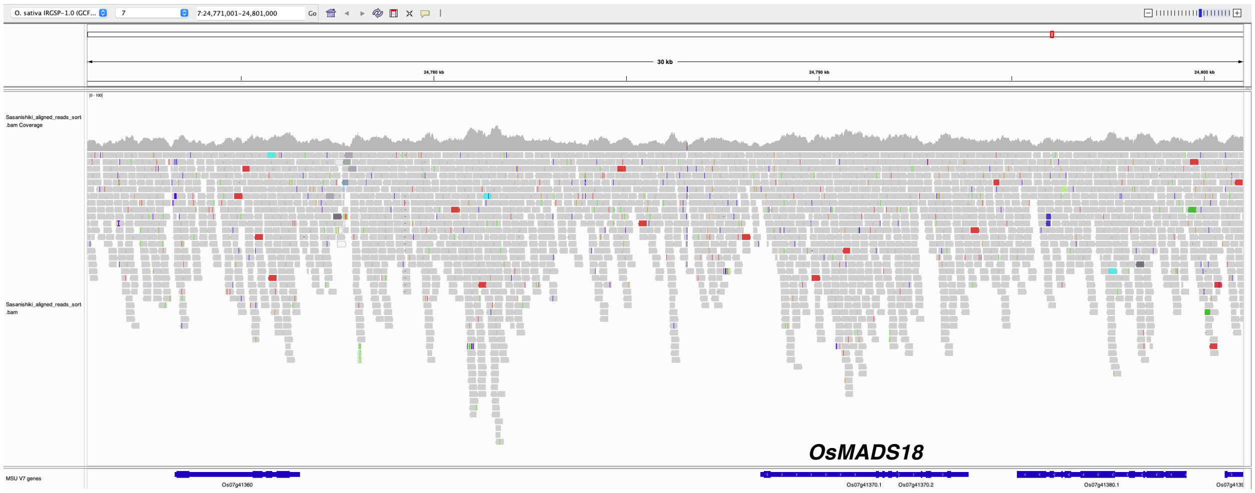


**(c)**

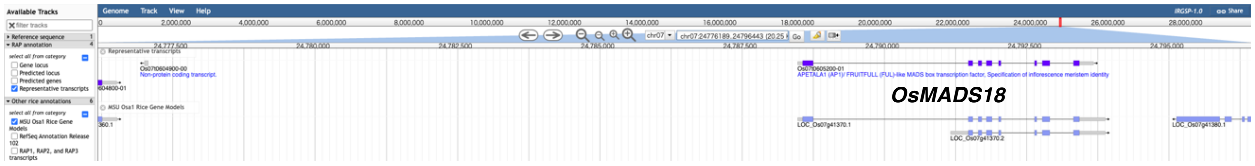

**Fig. S4** *OsMADS18* CNV in $T_0$ plants and progeny as determined by qPCR. (a) Frequency distribution of the relative amplification ratio for copy number determination in transgenic plants ($T_0$). (b) Summary table showing the numbers and percentages of $T_0$ plants with estimated copy number alterations. $T_0$ plants derived from different calli or from the same callus with different culture periods were counted as independent lines. The relative amplification ratio was used to classify heterozygous plants as follows: heterozygous for one copy/two copies, $0.3 < x < 0.7$; heterozygous for two copies/three copies, $1.3 < x < 1.7$; $x$ represents the relative amplification ratio. (c) Frequency distribution of progeny ($n = 400$) derived from plants estimated to be heterozygous for one copy/two copies (relative amplification ratio is ~0.5). (d) Frequency distribution of progeny ($n = 12$) derived from plants estimated to be heterozygous for two copies/three copies (relative amplification ratio is ~1.5).

(a)

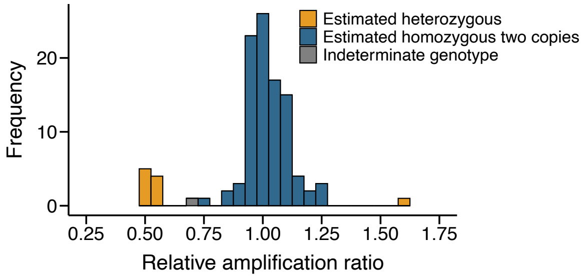


(b)

| | Culture period on selection medium | | Overall |
|---|---|---|---|
| | ~6 weeks | ~10 weeks | |
| Independently cultured $T_0$ plants | 76 | 31 | 107 |
| Estimated heterozygous one copy/two copies ($0.3 < x < 0.7$) | 5 (6.6%) | 4 (12.9%) | 9 (8.4%) |
| Estimated heterozygous two copies/three copies ($1.3 < x < 1.7$) | 0 (0.0%) | 1 (3.2%) | 1 (0.9%) |

(c)

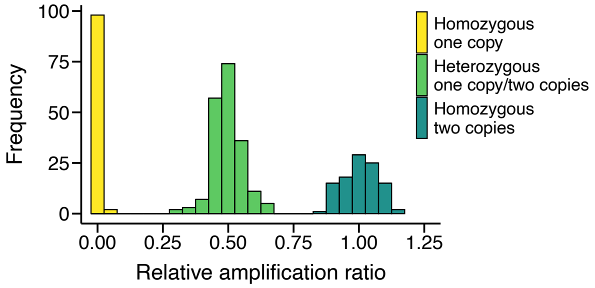


(d)

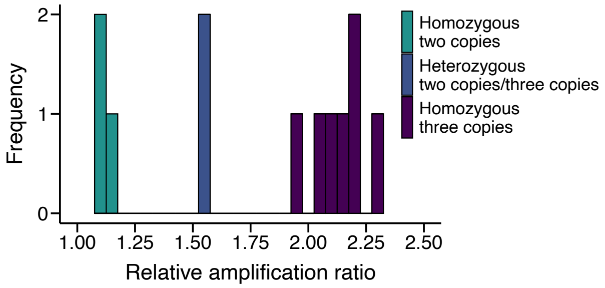

**Fig. S5** Genomic structure of the *OsMADS18* locus in IRGSP-1.0 and CNV-edited lines. (a, b) Dot matrix plots comparing the *OsMADS18* genomic region in the Nipponbare reference genome (IRGSP-1.0) to that in plants with one homozygous copy (1 CNV) (a) or three homozygous copies (3 CNV) (b) of *OsMADS18*. Grid lines are spaced at 0.03-Mb intervals. The orange shaded area marks the position of *OsMADS18*.

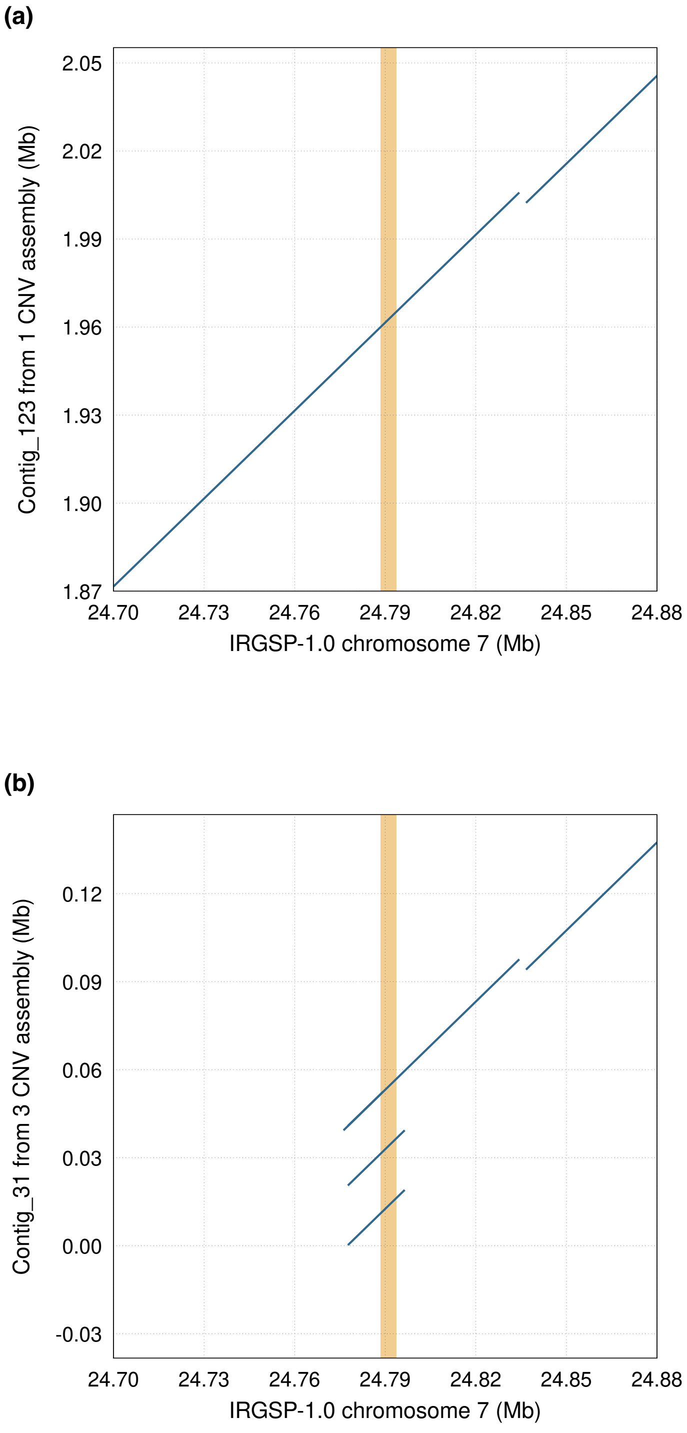

**Fig. S6** Effects of *OsMADS18* CNV on other agronomic traits. (a) Representative photograph of overall plant architecture. (b) Leaf blade length of the leaf just below the flag leaf ($n$ = 29, 35, and 29 for 1, 2, and 3 CNV, respectively). (c) Days to heading ($n$ = 30, 35, and 36 for 1, 2, and 3 CNV, respectively). "1 CNV," "2 CNV," and "3 CNV" refer to plants with one, two, and three homozygous copies of *OsMADS18*, respectively. Different lowercase letters indicate significant differences ($P < 0.05$) based on the Tukey-Kramer test for panels (b) and (c). All violin plots were generated using kernel density estimation with a smoothing adjustment factor of 1.5. All internal boxplots represent the first and third quartiles (the lower and upper hinges, respectively), the median (the middle horizontal line), and the range from the hinges to the smallest and largest observations within 1.5 times the interquartile range (the bottom and top whiskers, respectively). All data points, including outliers in the boxplot, are displayed as dots in the dot plot.

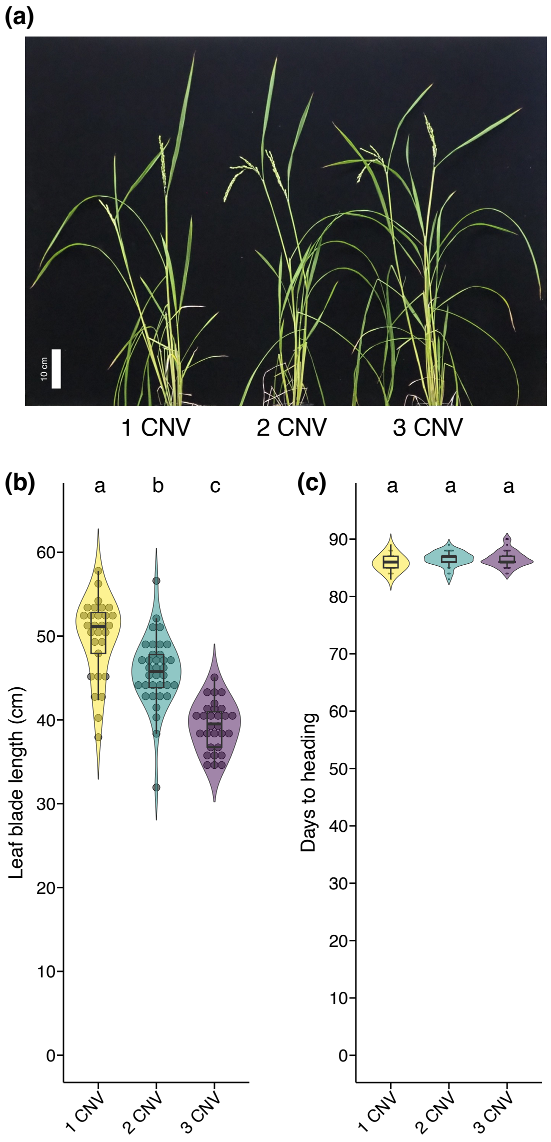

**Table S1** List of primers used in this study.

| Primer name | Sequence (5'-3') | Purpose/Target | Template | Experiment |
|---|---|---|---|---|
| GGsite-OsU6aPro-Target_F | TGAGAGGGTCTCTCTCGGGATTTTTTCCTGTAGTTTTCCCACAACCA | Cloning sgRNA cassette (*OsU6a* promoter) | pYLsgRNA-OsU6a | Plasmid construction |
| GGsite-OsU6aPro-Target_R | CCGCAACTTTACAGTACAGGCGGCAGCCAAGCCAGCA | Cloning sgRNA cassette (*OsU6a* promoter) | pYLsgRNA-OsU6a | Plasmid construction |
| GGsite-OsU6bPro-Target_F | AGTCAGGGTCTCTCTGATGCAAGAACGAACTAAGCCGGAC | Cloning sgRNA cassette (*OsU6b* promoter) | pYLsgRNA-OsU6b | Plasmid construction |
| GGsite-OsU6bPro-Target_R | CCGCAACTTTACAGTACAGGCAACACAAGCGGCAGCGC | Cloning sgRNA cassette (*OsU6b* promoter) | pYLsgRNA-OsU6b | Plasmid construction |
| Target-sgRNAScf-GGsite_F | CCTGTACTGTAAAGTTGCGGGTTTTAGAGCTAGAAATAGCAAGTTAAAATAAGGCT | Cloning sgRNA cassette | pYLsgRNA-OsU6a or pYLsgRNA-OsU6b | Plasmid construction |
| Target-sgRNAScf-GGsite_R1 | GCTGAAGGTCTCTTCAGTCCTTTGCTGCCGATTCCACAG | Cloning sgRNA cassette (*OsU6a* promoter) | pYLsgRNA-OsU6a or pYLsgRNA-OsU6b | Plasmid construction |
| Target-sgRNAScf-GGsite_R2 | GAGGTAGGTCTCTACCGTCCTTTGCTGCCGATTCCACAG | Cloning sgRNA cassette (*OsU6b* promoter) | pYLsgRNA-OsU6a or pYLsgRNA-OsU6b | Plasmid construction |
| pYL_Check_F | GAGCACCGGTAAGGC | Verification of the insert sequence | pYLCRISPR/Cas9$P_{ubi}$-H vector | Plasmid construction |
| pYL_Check_R | GCGCGCCAATGATACC | Verification of the insert sequence | pYLCRISPR/Cas9$P_{ubi}$-H vector | Plasmid construction |
| OsMADS18_Dup_Ctrl_F | GGTTGCTAAAGACCCATCAGG | The outside of the duplicated region | Genomic DNA | qPCR-based analysis of target gene copy number |
| OsMADS18_Dup_Ctrl_R | TACCCACGGACAAAATCAGC | The outside of the duplicated region | Genomic DNA | qPCR-based analysis of target gene copy number |
| OsMADS18_Dup_Junc_F | GTTCTTACCGTGCTCATCTTTGC | The junction of the duplicated region | Genomic DNA | qPCR-based analysis of target gene copy number |
| OsMADS18_Dup_Junc_R | CTTGGCAGTGCCGTTTGC | The junction of the duplicated region | Genomic DNA | qPCR-based analysis of target gene copy number |
| OsMADS18_F | TACGAGTTCTCCAGCCACTCC | *OsMADS18* | First-strand cDNA | RT-qPCR analysis |
| OsMADS18_R | ATTTGGCTCCAGTACGGCTCT | *OsMADS18* | First-strand cDNA | RT-qPCR analysis |
| UBQ5_F | ACCACTTCGACCGCCACTACT | *UBQ5* | First-strand cDNA | RT-qPCR analysis |
| UBQ5_R | ACGCCTAAGCCTGCTGGTT | *UBQ5* | First-strand cDNA | RT-qPCR analysis |

**Table S2** List of sequencing data and corresponding Sequence Read Archive (SRA) accession numbers.

| Sample name | Purpose | Sequencing platform | BioProject ID | SRA ID |
|---|---|---|---|---|
| Hitomebore | *De novo* genome assembly | Nanopore | PRJDB13864 | |
| Hitomebore | *De novo* genome assembly | Illumina | PRJDB13864 | DRR391466, DRR391467, DRR391468, DRR391469, DRR391470 |
| Hitomebore | Read depth-based CNV analysis | Illumina | PRJDB13864 | DRR391469, DRR391470 |
| Sasanishiki | *De novo* genome assembly | Nanopore | PRJDB20676 | |
| Sasanishiki | *De novo* genome assembly | Illumina | PRJDB20676 | |
| Sasanishiki | Read depth-based CNV analysis | Illumina | PRJDB20676 | |
| 1CNV | *De novo* genome assembly | Nanopore | PRJDB20676 | |
| 3CNV | *De novo* genome assembly | Nanopore | PRJDB20676 | |
| 1CNV_Rep1 | Read depth-based CNV analysis | Illumina | PRJDB20676 | |
| 1CNV_Rep2 | Read depth-based CNV analysis | Illumina | PRJDB20676 | |
| 1CNV_Rep3 | Read depth-based CNV analysis | Illumina | PRJDB20676 | |
| 1CNV_Rep4 | Read depth-based CNV analysis | Illumina | PRJDB20676 | |
| 1CNV_Rep5 | Read depth-based CNV analysis | Illumina | PRJDB20676 | |
| 2CNV_Rep1 | Read depth-based CNV analysis | Illumina | PRJDB20676 | |
| 2CNV_Rep2 | Read depth-based CNV analysis | Illumina | PRJDB20676 | |
| 2CNV_Rep3 | Read depth-based CNV analysis | Illumina | PRJDB20676 | |
| 2CNV_Rep4 | Read depth-based CNV analysis | Illumina | PRJDB20676 | |
| 2CNV_Rep5 | Read depth-based CNV analysis | Illumina | PRJDB20676 | |
| 3CNV_Rep1 | Read depth-based CNV analysis | Illumina | PRJDB20676 | |
| 3CNV_Rep2 | Read depth-based CNV analysis | Illumina | PRJDB20676 | |
| 3CNV_Rep3 | Read depth-based CNV analysis | Illumina | PRJDB20676 | |
| 3CNV_Rep4 | Read depth-based CNV analysis | Illumina | PRJDB20676 | |
| 3CNV_Rep5 | Read depth-based CNV analysis | Illumina | PRJDB20676 | |
